# *Toxoplasma gondii GRA28* is required for placenta-specific induction of the regulatory chemokine CCL22 in human and mouse

**DOI:** 10.1101/2020.10.14.335802

**Authors:** Elizabeth N. Rudzki, Stephanie E. Ander, Rachel S. Coombs, Hisham S. Alrubaye, Leah F. Cabo, Matthew L. Blank, Nicolas Gutierrez-Melo, JP Dubey, Carolyn B. Coyne, Jon P. Boyle

## Abstract

*Toxoplasma gondii* is an intracellular protozoan pathogen of humans that can cross the placenta and result in adverse pregnancy outcomes and long-term birth defects. The mechanism used by *T. gondii* to cross the placenta are unknown but complex interactions with the host immune response are likely to play a role in dictating infection outcomes during pregnancy. Prior work showed that *T. gondii* infection dramatically and specifically increases the secretion of the immunomodulatory chemokine CCL22 in human placental cells during infection.. Given the important role of this chemokine during pregnancy, we hypothesized that CCL22 induction was driven by a specific *T. gondii*-secreted effector. Using a combination of bioinformatics and molecular genetics, we have now identified *T. gondii* GRA28 as the gene product required for CCL22 induction. GRA28 is secreted into the host cell where it localizes to the nucleus, and deletion of this gene results in reduced CCL22 placental cells as well as a human monocyte cell line. The impact of GRA28 on CCL22 production is also conserved in mouse immune and placental cells both *in vitro* and *in vivo*. Moreover, parasites lacking GRA28 are impaired in their ability to disseminate throughout the animal, suggesting a link between CCL22 induction and the ability of the parasite to cause disease. Overall these data demonstrate a clear function for GRA28 in altering the immunomodulatory landscape during infection of both placental and peripheral immune cells, and show a clear impact of this immunomodulation on infection outcome.

**AUTHOR SUMMARY:** *Toxoplasma gondii* is a globally ubiquitous pathogen that can cause severe disease in HIV/AIDS patients and can also cross the placenta and infect the developing fetus. We have found that placental and immune cells infected with *T. gondii* secrete significant amounts of a chemokine (called “CCL22”) that is critical for immune tolerance during pregnancy. In order to better understand whether this is a response by the host or a process that is driven by the parasite, we have identified a *T. gondii* gene that is absolutely required to induce CCL22 production in human cells, indicating that CCL22 production is a process driven almost entirely by the parasite rather than the host. Consistent with its role in immune tolerance, we also found that *T. gondii* parasites lacking this gene are less able to proliferate and disseminate throughout the host. Taken together these data illustrate a direct relationship between CCL22 levels in the infected host and a key parasite effector, and provide an interesting example of how *T. gondii* can directly modulate host signaling pathways in order to facilitate its growth and dissemination.

## INTRODUCTION

*Toxoplasma gondii* is an obligate intracellular parasite that is an important parasite of humans and other animals. While this pathogen is particularly well-known to cause severe disease in the immunocompromised, such as those with HIV/AIDS or undergoing immunosuppression for organ transplants, *T. gondii* is also capable of crossing the placenta and infecting the developing fetus, leading to a variety of infection outcomes ranging from asymptomatic to severe (1). Importantly, even children born without symptoms are at high risk for extensive health problems later in life, including ocular disease and neurological disorders (2, 3). To date little is known about how *T. gondii* gains access to the fetal compartment and how the host responds to the presence of parasites at the maternal-fetal interface.

Recently we (4) found that primary human trophoblast cells (derived from term placentas) and 2nd trimester placental explants produced the chemokine CCL22 in response to infection with *T. gondii* (4). Production of this chemokine was dependent on parasite invasion and the dense granule effector trafficking gene product MYR1 (4). While the role of CCL22 during infection with *T. gondii* is poorly understood, this chemokine is a key molecular signal for the recruitment of regulatory T cells which are well known for their role in suppressing immune responses to tumors, leading to poor clinical outcomes (5, 6). Importantly disruption of T_reg_ recruitment to tumors can lead to improved outcomes in animal models. For example, using *Ccl22* DNA vaccines in mice leads to misdirection of regulatory T cells and ultimately reduced tumor growth (5). The role for CCL22 in healthy humans is less well understood, although it is thought to subvert and/or modulate inflammatory responses and may be particularly important for response resolution after pathogen clearance. CCL22 and regulatory T cells also play a critical role during pregnancy, where they seem to govern immune tolerance (7) and regulation of inflammation at the maternal-fetal interface. This regulatory role appears to be critical in determining pregnancy outcome during pathogen-mediated immune activation (7, 8). Given the important role played by CCL22 during pregnancy and our recent findings regarding the ability of a congenitally acquired parasite to directly modulate production of this chemokine, we sought to identify the parasite effector(s) responsible for this in order to determine the impact of CCL22 modulation on congenital transmission and pregnancy outcome during vertical transmission. To do this we used a bioinformatic screen identify candidate genes and identified one (TGGT1_231960) as being required for CCL22 induction in human and mouse cells. Overall these data show that a specific effector is largely responsible for *T. gondii*-mediated CCL22 induction in a relatively small number of human and mouse cell types, and suggest that the manipulation of CCL22 levels by GRA28 may influence the ability of *T. gondii* to disseminate to throughout the host.

## RESULTS

### *Toxoplasma gondii* induces a monocyte-like cell line to produce the immunomodulatory chemokine CCL22

Previous work established that placental explants and primary human trophoblasts infected with *T. gondii* had increased *CCL22* transcript abundance and released more CCL22 protein into the culture media compared to mock infected controls (4). Since we also found that not all cell types produce CCL22 in response to infection (e.g., HFFs), we were interested in identifying a human cell line that could be used as a more tractable model than placental cells to assay *T. gondii*- driven CCL22 induction. THP-1 cells, a cell line derived from a patient with monocytic leukemia, were a reasonable candidate given their origins in the myeloid lineage and known production of CCL22 in response to a variety of stimuli (9, 10). We infected THP-1 cells with a type 1 *T. gondii* strain (RH88 or RH:YFP;(4)) at a multiplicity of infection (MOI) of 3. Following 24 hours of infection, supernatants were collected from each well. Human foreskin fibroblasts (HFFs) were infected in parallel as negative controls. Mock treatments involved passing the parasite solution through a 0.22 µM filter prior to exposure to the cells. Based on CCL22 ELISA, *T. gondii* infection induced CCL22 in THP-1 cells, and as expected there was no CCL22 production from mock-treated controls or *T. gondii*-infected HFFs (**Figure 1a**). We also infected primary placental tissues in the same manner, and as expected villous tree explants and decidua taken from 2nd trimester placentas produced significantly more CCL22 compared to mock-treated controls (**Figure 1a**). In addition to the type 1 RH strain, other *T. gondii* strain types (Type 2:PRU and Type 3:CEP; Supplementary **Figure 1 Supplement 1a**) also induced secretion of CCL22 from THP-1 cells, as did the nearest extant relative of *T. gondii*, *Hammondia hammondi* (**Figure 1 Supplement 1b**). In contrast to *H. hammondi*, and just as we observed previously in primary human placental cells (4), *Neospora caninum* has no effect on THP-1 production of CCL22 (Supplementary **Figure 1 Supplement 1b**). These data provided strong support that the mechanism of CCL22 induction was the same for THP-1 and placental cells.

**Figure 1:**
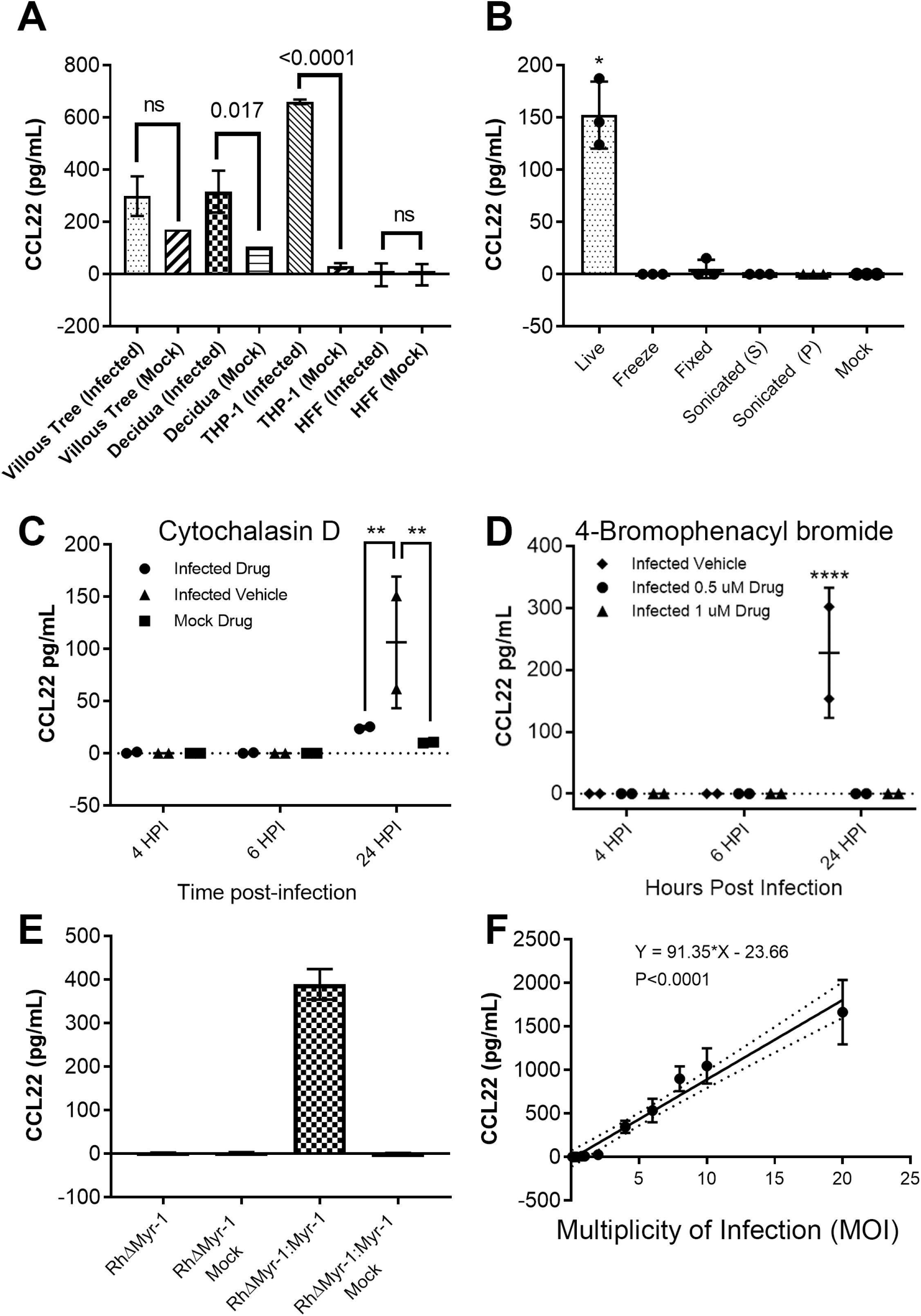
A) THP-1 cells, Human Foreskin Fibroblasts (HFFs), and 2nd trimester placental samples of villous trees and decidual tissue were infected with a type I strain of *Toxoplasma gondii* (RH:YFP). Statistics on the THP-1 cell and HFF cell samples performed by ordinary Two-way ANOVA analysis with Tukey’s multiple comparisons test. Statistics on the placental samples performed by two-tailed welch-corrected t-tests. B) Type I strain (RH) *T. gondii* parasites subjected to multiple treatments as described in methods. The soluble fraction of the sonicated treatment is denoted by S, the insoluble fraction is denoted by P. THP-1 cells were then exposed to either live parasites, or treated parasites. Statistics performed by multiple two-tailed welch-corrected t-test comparisons with the live parasite treatment *P* ≤ 0.0145. C,D) Type I strain (RH) *T. gondii* parasites were treated with either Cytochalasin D (C), or 4- Bromophenacyl bromide (D) as described in methods. THP-1 cells were then infected with the respective parasite treatment. Statistics were performed by two-way ANOVA analysis and multiple comparisons post-hoc tests, where Cyt-D *P* = 0.009, and 4-BPB *P* < 0.0001. E) Type I strain (RH) *T. gondii* parasites deficient in Myr-1 (TgRHΔMyr-1) and their complement (TgRHΔMyr-1:Myr-1_comp_) were used to infect THP-1 cells. (F) Type I strain (RH) *T. gondii* parasites were used to infect THP-1 cells at MOIs of: 20, 10, 8, 6, 4, 3, 1, 0.8, 0.4, 0.2, and 0.1. A, B, C, D) Respective cells/tissues were infected with a multiplicity of infection (MOI) of 3. Supernatants were collected at 24 hours post-infection unless indicated otherwise, and assayed by CCL22 ELISA.

We also determined if live parasites were required to induce CCL22 in THP-1 cells by exposing host cells to parasites that were exposed to a variety of lethal treatments. As shown in **Figure 1b** dead parasites failed to induce CCL22 production by THP-1 cells. We also pretreated parasites and host cells with 10 µg/mL Cytochalasin-D (Cyt-D) to block invasion (11), and as shown in **Figure 1c** Cyt-D treated parasites were significantly impaired in their ability to induce CCL22, suggesting that active invasion was required for this phenomenon. We obtained similar results with the inhibitor 4-BPB (**Figure 1d**), which also significantly blocked CCL22 production by THP-1 cells at 0.5 and 1 µM. This drug blocks rhoptry and dense granule secretion from *T. gondii*, but not microneme secretion(12), suggesting that the factor is not a microneme protein (**Figure 1d**). We also infected THP-1 cells with *T. gondii* parasites that were deficient in the dense granule trafficking protein MYR1 ((13); kind gift from John Boothroyd, Stanford University) and compared them to TgRH*ΔMYR1*:MYR1_c_ parasites. TgRH*ΔMYR1* parasites failed to induce any detectable CCL22 from THP-1 cells while, as expected, TgRH*ΔMYR1*:MYR1_c_ parasites induced significantly more than mock-treated cells (**Figure 1e**). We also observed a very tight correlation between parasite multiplicity of infection (MOI) and CCL22 levels, suggesting that the signal was primarily driven by the parasite rather than the host cell (**Figure 1f**). Based on these results we felt confident that the unknown secreted factor driving CCL22 production in human primary placental cells was very likely the same as the one driving it in the THP-1 cell line and chose THP-1 cells for screening candidate effectors given their tractability in the laboratory.

### Transcript abundance correlation analysis identifies a large group of putatively MYR1-trafficked gene products

As described previously (above and (4)), we have determined that primary human trophoblast cells infected with *T. gondii* have a transcriptional signature that is characterized by the production of immunomodulatory chemokines, with CCL22 being the most potently induced. To identify candidate *T. gondii* genes responsible for this effect on placental cells, and since this effect required the *T. gondii* effector translocation complex protein MYR1 (4), we hypothesized that MYR1-dependent substrates would have highly correlated gene expression profiles across diverse gene expression datasets. To test this hypothesis, we generated an “all vs. all” correlation matrix of 396 *T. gondii* Affymetrix microarray datasets that we downloaded and curated from the Gene Expression Omnibus (see Materials and Methods). Analysis of the entire correlation matrix (shown in **Figure 2a** and downloadable at 10.6084/m9.figshare.16451832) confirms this hypothesis for certain gene classes. For example, we identified one cluster containing multiple SAG-related sequences (SRS) which are typically expressed at high levels in bradyzoites (including SRS49 cluster members A, C and D; **Figure 2 Supplement 1A**) and another containing 70 genes, 43 of which encode ribosomal subunits (**Figure 2 Supplement 1a**). Examination of the gene expression heatmaps across all 396 microarray analyses clearly show distinct patterns of gene expression in these two clusters depending on life stage treatment exposure (**Figure 2 Supplement 1a,b**).

**Figure 2:**
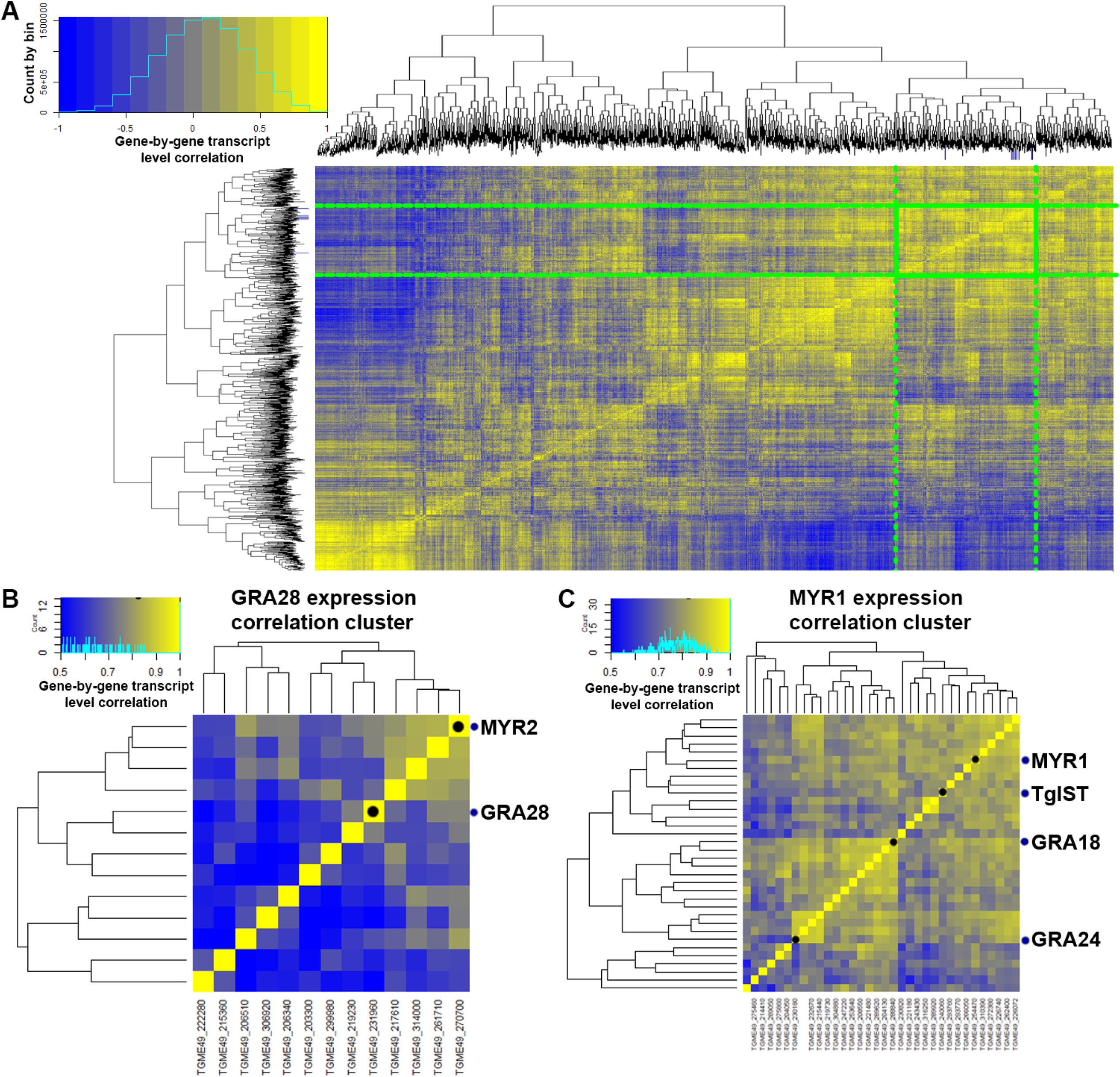
Transcript correlation analyses to identify putative MYR1 substrates based on 396 publicly available microarray datasets. A) Gene-by-gene correlation data for *T. gondii* genes across 396 microarray datasets. A subset (3,217 genes with at least one sample having a normalized log2-transformed value ≥10) of the total number (8,058) of *T. gondii* genes are shown for simplicity. Genes outlined in the green box indicate the cluster containing all of the bait genes as well as candidate CCL22-inducing genes with the exception of *Toxofilin*. Dark blue tick marks on each dendrogram indicate the location of all of the bait genes. Color scale covers correlations ranging from -1 to +1. B) Subcluster containing *MYR2* and *GRA28*. C) Subcluster containing *MYR1*, *TgIST*, *GRA18* and *GRA24*. Note: For B and C the color scale is from 0.5 to 1.0 to highlight subcluster differences.

We quantified the degree of transcript abundance correlation between 5 “bait” genes (*MYR1*, *2* and *3*; (13, 14)) and the known *MYR1*-dependent substrates *TgGRA24* and *TgIST* (15, 16)) and all other genes across all 396 expression datasets. We identified genes as candidate MYR1 substrates if they had an average correlation with the 5 “bait” genes ≥0.7, a dN/dS ratio ≥2, and the presence of a predicted signal peptide OR at least one transmembrane domain. Using this set of filters we were left with 28 candidate genes (plus all 5 bait genes which also met these cutoffs), including the known TgMYR-dependent substrate TgGRA25 (17). Since all known MYR1 trafficked substrates are dense granule proteins, we eliminated any surface antigens or soluble enzymes, leaving a number of confirmed dense granule proteins (e.g., GRA4 and GRA8) and conserved hypothetical proteins. Importantly, when we examined the correlation between the bait genes and all *T. gondii* genes annotated as “Dense granule” either in the primary product notes or via user annotation, we found that not all dense granule- encoding transcripts correlated highly with bait transcript levels (**Figure 3 Supplement 1a**), indicating that our approach could discriminate between different classes of proteins secreted from the same organelle. For example, while genes like *GRA32* (*TGME49_212300*) had transcript levels with relatively high (>0.8) correlations with bait transcript levels, other genes encoding GRA1, GRA2 and GRA11 paralogs had transcript levels that correlated much more poorly with the bait genes. This is despite the fact that most of these dense granule encoding genes have high transcript levels compared to other gene clusters as shown in the heat map in **Figure 3 Supplement 1a,** demonstrating that our approach yielded an additional layer of discrimination to categorize dense granule-trafficked gene products. Moreover, many of the co-regulated genes are not yet annotated but based on our analysis one would predict that many are likely to be dense granule protein derived secreted effectors or structural constituents of this parasite organelle.

### *T. gondii GRA28* is the gene responsible for CCL22 induction in human immune and placental cells

When we specifically examined correlations between the bait genes listed above and *MYR1*, we found that *MYR1* expression profiles were highly correlated at the transcriptional level with *MYR2/3* and *IST* (**Figure 2b,c** and **Figure 3a**, top), consistent with the idea that MYR1 substrates could be identified using this approach. After identifying a small list of candidate genes (**Figure 3a**, bottom), we deleted each using CRISPR-CAS9 and screened for CCL22- induction in THP-1 monocytes by ELISA. Among the five genes that we tested (including *GRA18* which was recently found to induce Ccl22 in mouse macrophages; (18)), we found that *GRA28* (*TGME49_231960*) was required for the induction of CCL22 secretion by infected THP- 1 monocytes (**Figure 3b**). We also found that Δ*Toxofilin* parasites had significantly reduced levels of CCL22 induction (**Figure 3b**), albeit to a much lesser extent than *ΔGRA28* parasites. We think it likely that this decrease is owed to the reduced invasion capacity of Δ*Toxofilin* parasites rather than a direct impact of this gene product on host CCL22 production (19, 20).

**Figure 3:**
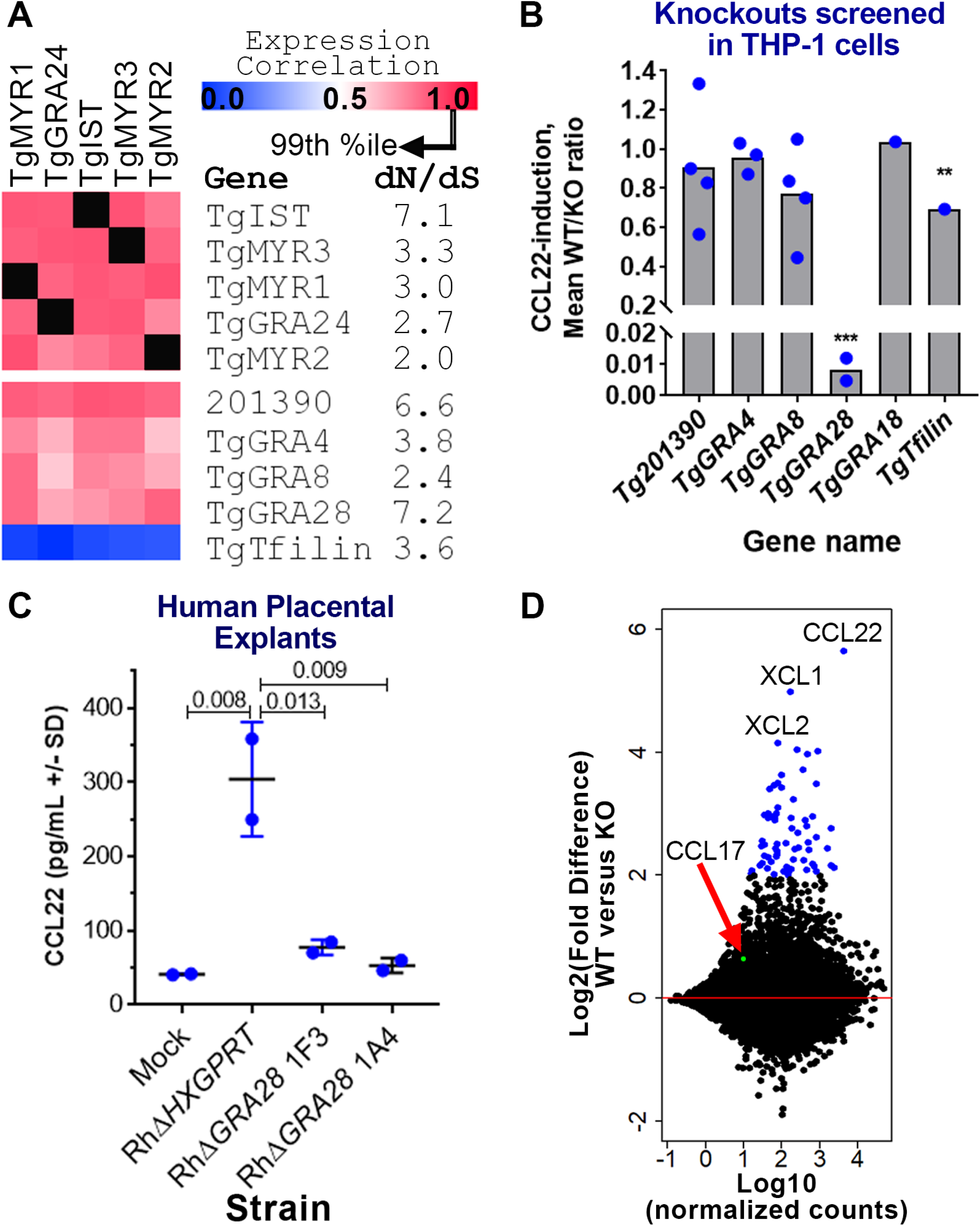
Identification of the *T. gondii* gene GRA28 as an inducer of CCL22 in human cells. A) Gene expression correlations across 396 *T. gondii* expression microarrays between TgMYR1 and 4 additional “bait” genes (top) and 5 candidate CCL22-inducing effectors (bottom). dN/dS ratios are also shown to illustrate the high level of positive selection acting on this class of genes. B) Effect of deleting 5 candidate genes on CCL22 secretion in THP-1 cells, showing that *ΔGRA28* parasites induced significantly less CCL22 compared to wild type controls (>100 fold reduction; ***P<0.0001). PRUΔ*Toxofilin* parasites also induced significantly lower levels of CCL22 in THP-1 cells (1.4-fold reduction; **P<0.01). Each blue dot indicates a genetically distinct knockout clone. C) *ΔGRA28* parasite clones also induce significantly less CCL22 from primary human 2nd trimester placental villous explants. D) MA plot of RNAseq analysis performed on THP-1 cells infected with WT or *ΔGRA28 T. gondii* (RH strain). CCL22 and the chemokines CXL1 and CXL2 were the most highly GRA28-dependent transcripts, while a handful (64) of other genes had significantly higher transcript abundance in WT parasites compared to *ΔGRA28* parasites (Padj<0.001; Log2FC>2; blue symbols). CCL17 (arrow, green symbol), a chemokine that is typically co-regulated with CCL22, did not show any evidence of being induced by GRA28.

To determine if GRA28 was responsible for CCL22 production by human placental cells, we infected 2nd trimester human villous placental explants with WT and ΔGRA28 parasites and observed a marked decrease in CCL22 production by explants exposed to *ΔGRA28* parasites compared to WT (**Figure 3c**). To gain a broader understanding of the transcriptional networks altered by *GRA28* we compared THP-1 cells infected with RHΔ*HPT*:*HPT* and RH*ΔGRA28* using RNAseq. A relatively small number of transcripts had significantly altered abundance using stringent statistical cutoffs (67 genes with Padj<0.0001 and abs(log_2_FC≥2) are highlighted in **Figure 3d**) and these included CCL22 as well as the chemokines XCL1 and XCL2. Interestingly transcript abundance for CCL17, a chemokine that is often co-regulated with CCL22 (21, 22) and which is induced in some cells along with CCL22 by the *T. gondii* effector GRA18 (18), was not dependent on *GRA28* (**Figure 3d**). The majority of transcripts that were GRA28-dependent were of higher abundance in WT compared to *ΔGRA28* parasites when using slightly relaxed statistical cutoffs (263 higher, 33 lower; P_adj_<0.05 and log_2_FC≥1 or ≤1).

We performed pathway analysis on these sets of regulated genes using Ingenuity pathway analysis (IPA) and identified host cell pathways that were either more or less induced in WT *T. gondii*-infected cells compared to *ΔGRA28*-infected cells (**Figure 4**), including <u>Dendritic Cell Maturation</u>, <u>IL6 and IL8 signaling</u>, and <u>NFκB signaling</u> (**Figure 4a**). When we assessed the degree of gene overlap in these gene sets, we found that 10 of the pathways contained the *JUN* and *FOS* genes (**Figure 4b**), indicating a potential role for AP-1 complex targeted transcripts in GRA28-dependent transcriptional changes. We examined correlations across these gene sets (after creating a matrix of presence/absence of each of the genes shown in **Fig. 4b**) and identified two non-overlapping sets of genes. The larger cluster contains multiple immunity-related genes while the smaller cluster contains genes involved in proteoglycan synthesis (**Figure 4c**), including the *XYLT1* gene which encodes the enzyme that adds UDP-Xylose to serine residues as a first step in glycosaminoglycan synthesis. When we performed a similar analysis using the “upstream regulator” module in IPA, we identified a small set of significant (Z-score ≥2; *P*<0.001) regulatory factors that were upstream of the GRA28- dependent gene set, including multiple regulators associated with the NFκB pathway (**Figure 4 Supplement 1a**). Cluster (**Figure 4 Supplement 1b**) and downstream gene overlap (**Figure 4 Supplement 1c**) analyses further confirmed the *FOS* and *JUN* genes as contributing to the signaling pathways that were GRA28-dependent, while also confirming a putative role for NFκB. For example, the cluster with the most similar target gene overlap contains multiple genes in the NFκB pathway (*NFKBIA*, *NFKB1*, *RELA*) (**Figure 4 Supplement 1c**). However, when we co-transfected HEK293 cells with NFκB luciferase reporter constructs and a construct encoding the first exon of *GRA28* (see below) we saw no increase in the levels of luciferase after GRA28 transfection in contrast to a known NFκB activating construct containing multiple Caspase Activation and Recruitment Domains (CARDs; **Figure 4d**). This suggests that NFκB activation may not play a role in CCL22 induction. Other candidate transcriptional mediators with GRA28- dependent transcript levels are *FOS*, *JUN* and *IRF4* (**Figure 4 Supplement 1d**). Transcript levels of *JUN* have been shown in numerous studies to increase in a variety of host cells after infection with *T. gondii* (23, 24). To test whether GRA28 played a role in altering C-JUN abundance during infection we infected THP-1 cells with WT or ΔGRA28 *T. gondii* parasites for 24 h and using semi-quantitative western blotting to quantify C-JUN protein levels. While infection of THP-1 cells clearly increased C-JUN levels compared to mock-treated cells (**Figure 4 Supplement 2**), the presence or absence of GRA28 in the infecting strain had no significant impact on C-JUN protein abundance. These data suggest that while JUN transcript levels appear to be at least somewhat dependent on GRA28 in the infecting strain (**Figure 4 Supplement 1d**), this does not appear to be detectable at the protein level using western blotting.

**Figure 4.**
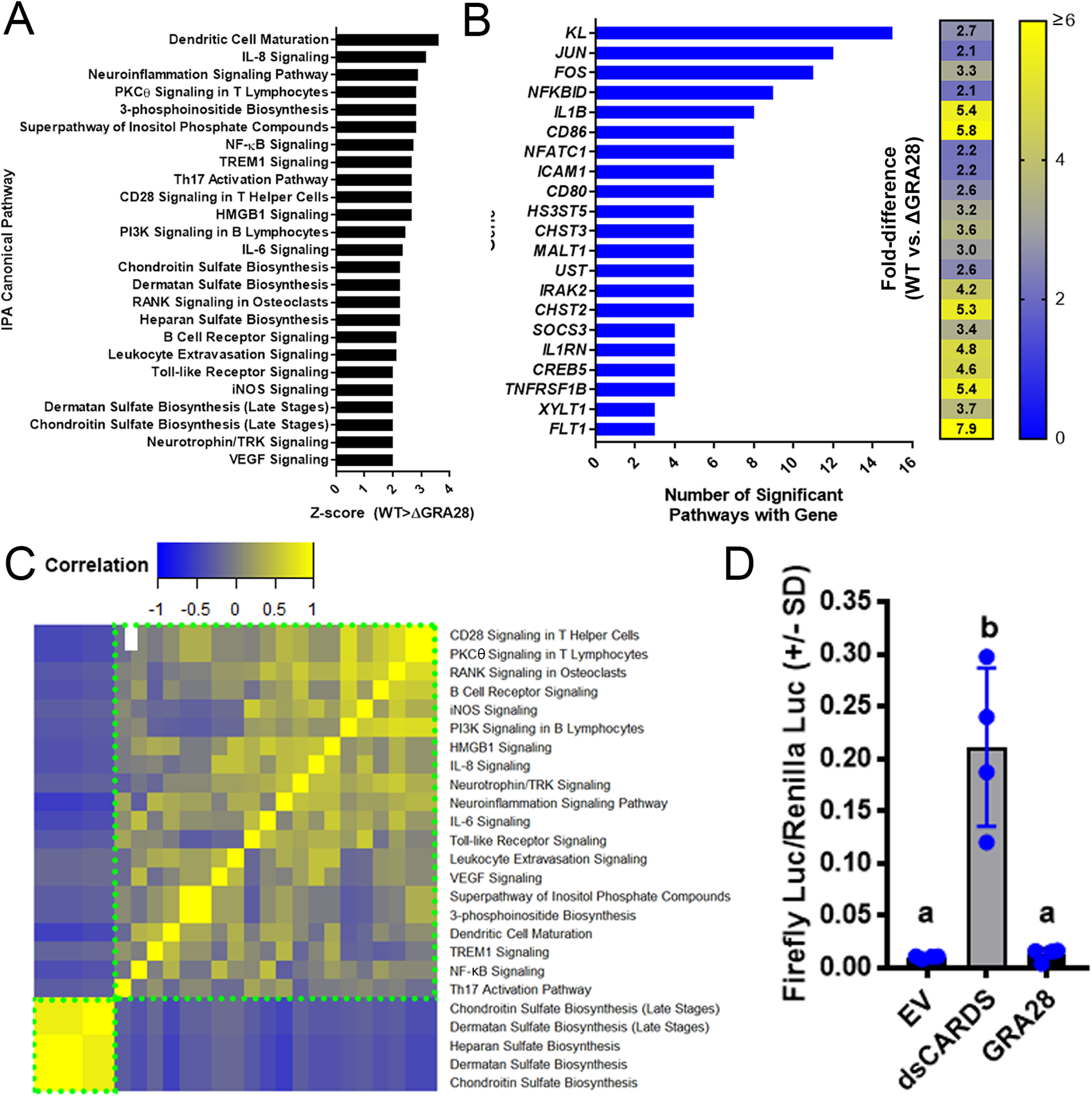
IPA analysis on THP-1 cells infected with WT or *ΔGRA28 T. gondii* parasites for 24h showing canonical pathways that were differentially regulated (-log(*P*)≥2; Z score ≤ -2 or ≥ 2) depending on the presence or absence of the GRA28 gene. A) Z scores for significant canonical pathways. All were higher in WT compared to *ΔGRA28 T. gondii*. B) There was extensive overlap of component genes within each canonical pathway, particularly for the genes *KL* and those encoding components of the AP-1 transcription factor complex (*JUN* and *FOS*). A heatmap of fold-difference in transcript abundance for cells infected with RH:WT and RH*ΔGRA28* is shown. C) GRA28 is responsible for driving transcriptional changes in two major gene clusters identified based on the degree of gene sharing between each canonical pathway (clusters outlined in dotted green boxes). The larger cluster consists primarily of immunity- related pathways while the smaller cluster consists of genes involved in proteoglycan synthesis. D) Quantification of NFκB activation in 293T cells. Cells were transfected with NFκB firefly luciferase plasmid, a consitutive renilla luciferase plasmid, as well as empty vector (EV), a construct expressing a CARD domain (dsCARDS) or the first exon of *T. gondii* GRA28. While the CARD domain construct induced firefly luciferase expression as expected, expression of *T. gondii* GRA28 had no significant impact on firefly luciferase levels (letters indicate groups that were not significantly different from one another following one way ANOVA and Tukey’s multiple comparisons post-hoc test.)

### The first exon of *GRA28* is sufficient for induction of CCL22 in during parasite infection of and ectopic expression in human cells

The *GRA28* gene has been described previously as encoding a dense granule protein that was capable of trafficking to the host cell nucleus during infection (25). However, the exact structure of the GRA28-encoding gene was somewhat ambiguous based on its annotation in ToxoDB. Specifically, while TGME49_231960 is predicted as a single exon gene spanning ∼7.4 kb of genomic sequence (**Figure 5a**), the annotated gene is shorter in TGGT1 (**Figure 5a**) and split into two gene products in *T. gondii* strains VEG, FOU, ARI, VAND, MAS, CATPRC2 and P89. The 5’ end of the gene was consistently predicted across all annotated genes, including the precise location of the first intron. When we performed *de novo* assembly of the *T. gondii* RH transcriptome, we were unable to identify any assembled transcripts that spanned the entire length of the TGME49_231960 prediction, most likely due to the fact that a 39 bp repeat in between each of these transcripts disrupted the assembly process (repeat consensus sequence: CAGCAGCAGCCACAAGGGWMTGTTGTGCATCAACCACTA; **Figure 5a**). However it should be noted that when we examined recently released Oxford Nanopore long-read single molecule sequencing of *T. gondii* transcripts that are available on ToxoDB.org there are multiple reads that span this repeat region (select Nanopore reads shown in **Figure 5a**), suggesting that the gene is at least similar to that predicted for ME49 in the *Toxoplasma* genome database. Regardless, given the challenges associated with amplifying and cloning this repetitive region we expressed an HA-tagged version of the first exon of GRA28 in *T. gondii* and observed expression within both the parasites and HA signal in the nucleus of infected cells (**Figure 5b**). Importantly, CCL22 induction could be restored in an RH*ΔGRA28* clone after bulk transfection of the exon 1 GRA28-expression construct prior to infecting THP-1 cells (**Figure 5c**), confirming the role of sequences present in the first exon of GRA28 in driving CCL22 production in human cells. Similar results were obtained when we transiently expressed a construct containing the entire genomic locus for the predicted *T. gondii* GT1 *GRA28* gene (light green bar, **Figure 5a**) in RH*ΔGRA28* parasites (**Figure 5d**). In contrast, when the Exon 1 construct was expressed transiently in *Neospora caninum* (strain NC-1; (26)), we did not observe any HA signal in the infected host cell despite expression of the protein within the parasite (**Figure 5e**). When we quantified host nuclear HA signal intensity (background subtracted and then normalized to staining intensity within the parasite; see Materials and Methods) in infected host cells there was a clear and significant (*P*=0.0012) difference in the amount of HA-derived signal in the host nucleus when TgGRA28 was expressed in *T. gondii* compared to *N. caninum* (**Figure 5 Supplement 1a**). Close inspection of multiple images suggest that the trafficking of *T. gondii* GRA28 within *N. caninum* itself may be distinct from how it traffics in *T. gondii*. For example HA staining was observed mostly within the parasite for *N. caninum* but could be found both within *T. gondii* and at the vacuole periphery (**Figure 5e** and **Figure 5 Supplement 1**). While clearly *N. caninum* failed to traffic detectable amounts of GRA28 into the host cell, this could be due to a) poor trafficking of the protein within the parasite such that it never gains proper access to vacuolar export machinery components like MYR1 and/or b) poor trafficking from the parasite into the host cell due to incompatibility with the *N. caninum* export machinery. Interestingly *N. caninum* does not appear to have an intact *GRA28* gene in its genome (see the synteny map for TGME49_231960 at ToxoDB.org), although it does have a *MYR1* ortholog which has been shown to be sufficient to traffic secreted *T. gondii* proteins into the host nucleus. Finally, GRA28-Exon1 (minus the residues encoding the predicted signal peptide) could be robustly expressed in HeLa cells with a V5 tag where it trafficked to the host cell nucleus (**Figure 5f**) and also was functional when expressed ectopically in THP-1 cells where it induced CCL22 secretion (**Figure 5f**).

**Figure 5:**
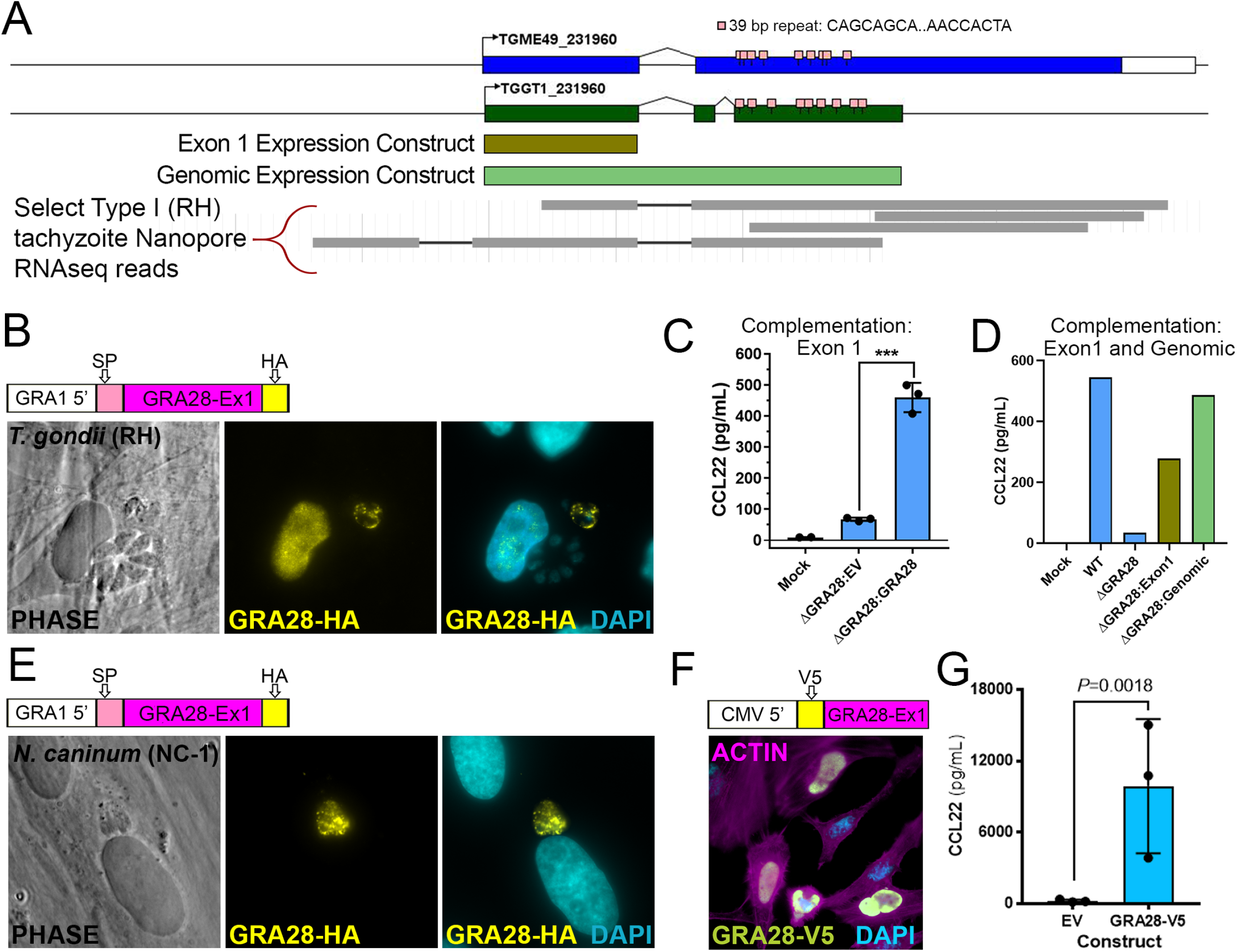
A) Schematic of the *GRA28* locus along with its gene prediction in the current annotation of the *T. gondii* genome (www.toxodb.org). In addition to gene models from two *T. gondii* strains (GT1 and ME49), 39 bp repeats and regions used in expression constructs are shown in brown and light green. Map created using GenePalette software (see Materials and Methods; (51)). B) Sequence encoding an N-terminal HA tag was inserted immediately after the predicted signal peptide cleavage site in a GRA1-promoter driven version of *T. gondii* GRA28 Exon 1. When transiently transfected into *T. gondii* HA-tagged protein could be detected in the parasites as well as the host cell nucleus. C) *ΔGRA28 T. gondii* parasites (RH strain) were transiently transfected with empty pGRA-HA-HPT vector (EV) or the same construct described in B encoding an HA-tagged version of *T. gondii* GRA28 Exon 1. After washing in cDMEM parasites were used to infect freshly plated THP-1 cells for 24h and CCL22-levels were quantified in culture supernatants using ELISA. Mock-treated cells were exposed to a sterile filtered parasite preparation. D) E) The construct encoding HA-tagged *T. gondii* GRA28 Exon 1 (same as that used in B and C) was used to transfect *Neospora caninum*, a near relative of *T. gondii*. HA staining revealed expression of this *T. gondii* GRA28 Exon 1 in *N. caninum* parasites (visualized by HA staining) but in contrast to *T. gondii* we did not observe trafficking of GRA28 to the host cell nucleus when expressed in this strain. Quantification of nuclear HA-derived signal is presented in Figure 5 Supplementary Figure 1. F) Sequences encoding an N-terminal V5 tag were inserted downstream of a Kozak consensus sequence and upstream of GRA28 Exon 1 (minus the signal peptide-encoding sequence). The construct was transfected into HeLa cells and V5 staining was observed prominently in the nucleus of transfected cells. G) Transfection of the construct in E directly into RAW 264.7 cells significantly induced CCL22 production as detected by ELISA. T-test was performed on log_10_-transformed data.

### GRA28 induction of Ccl22 is fully conserved in mice

To determine whether parasite-driven induction of CCL22 is conserved in the murine model, we compared WT and GRA28-deficient (*ΔGRA28*) parasites for their ability to induce this chemokine *in vitro*, *ex vivo*, and *in vivo*. First, we infected mouse macrophages (RAW 264.7) *in vitro* with type 1 strain (RH) *T. gondii* parasites (WT), or (RH) *ΔGRA28 T. gondii* parasites at MOIs of 3. Based on Ccl22 ELISA, mouse macrophages not only release more Ccl22 protein during *T. gondii* infection, but similar to human THP-1 cells this phenotype is also dependent on the presence of *T. gondii* secreted protein GRA28 (**Figure 6a**). Next, we investigated whether primary mouse tissues, specifically mouse placental tissue, also elicit this response to *T. gondii* infection. Embryonic day 12.5 Swiss Webster mouse placentas were halved and distributed into separate treatment groups. These placental explants were then infected *ex vivo* with 2.0 x 10^6^ Type 1 strain (RH) *T. gondii* parasites (WT), (RH) *ΔGRA28 T. gondii* parasites, or mock treatment. As shown in **Figure 6b**, primary mouse placental tissue also responds to *T. gondii* infection by releasing Ccl22 protein in a GRA28-dependent manner. RNA was also extracted from the infected placental samples and we performed RNAseq. As shown in **Figure 6c** the number of transcripts that varied in a GRA28-dependent manner was markedly small, suggesting that GRA28 is a highly specific inducer of *Ccl22* in mouse placental explants. Of the three genes with significantly higher transcript levels (*Ccl22*, *Il12rb2*, *Ccr7*) in wild type 1nfections as compared to *ΔGRA28* infections, *Ccl22* was the most highly induced. These data show conservation of the parasite-driven Ccl22 phenotype 1n primary mouse placental explants at both a protein and transcript level. Finally, we investigated mouse *in vivo* Ccl22 responses to *T. gondii* intraperitoneal infection. Female BALB/cJ mice (n = 3 for each treatment) were infected with WT, *ΔGRA28*, or mock *T. gondii* treatments. We focused on early, acute infection and performed Ccl22 ELISA on serum (**Figure 6d**) and peritoneal lavage fluid (**Figure 6e**). These suggest *in vivo* Ccl22 protein levels are at least partially dependent on GRA28. Moreover, while there was a significant amount of systemic Ccl22 protein detected in serum of infected mice, even in the *ΔGRA28* parasite treatment, Ccl22 was almost undetectable in peritoneal lavage fluid in *ΔGRA28*-infected mice. Overall, these data indicate the process driving *T. gondii* GRA28 induced Ccl22 is similar, if not the same, in both mice and humans, and that this parasite effector can mediate robust changes in Ccl22 production at the site of infection and systemically.

**Figure 6.**
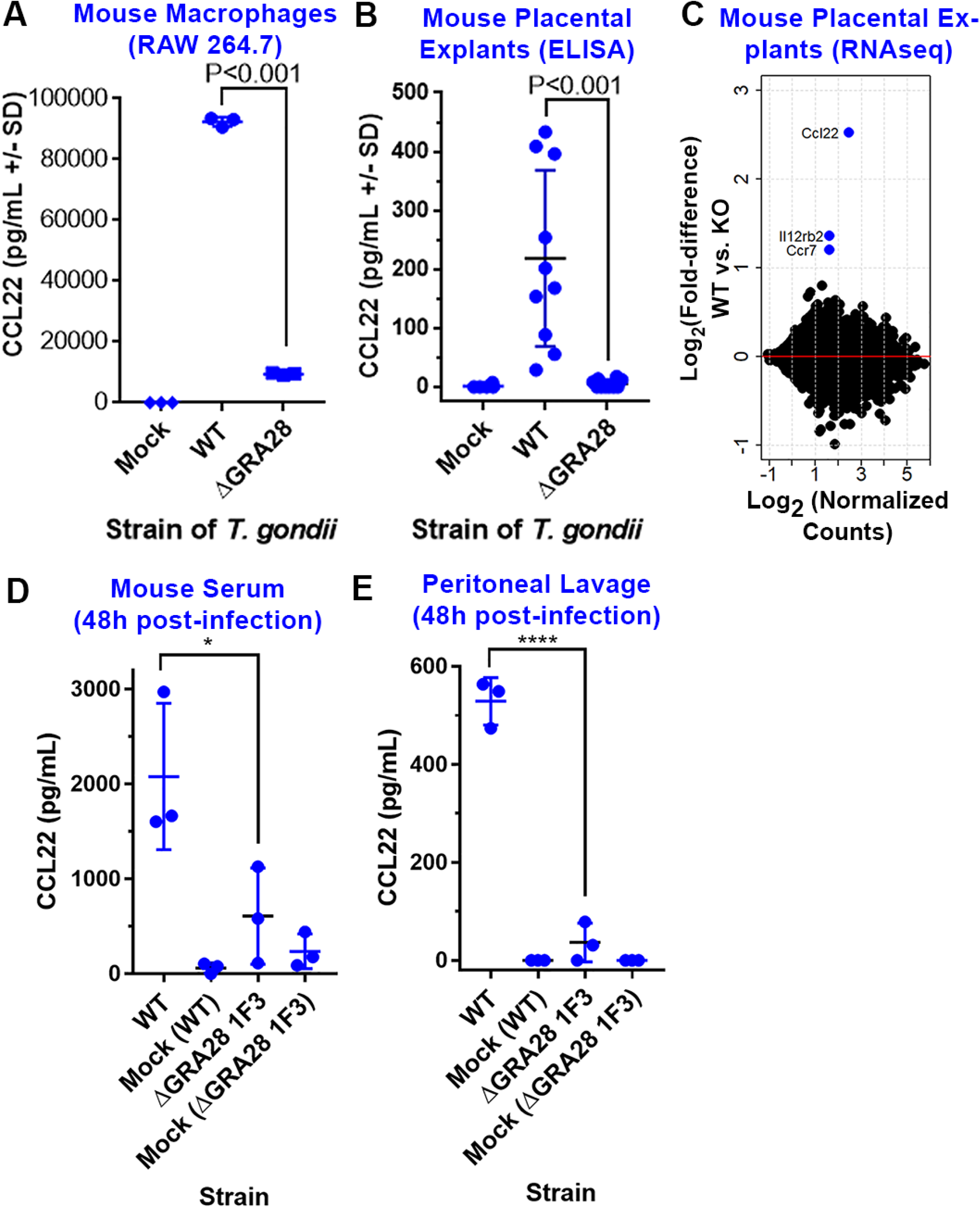
GRA28 induction of Ccl22 is conserved in mouse immune and placental tissues. A,B) *ΔGRA28* parasites induce significantly less Ccl22 secretion from RAW 264.7 macrophages (A) and mouse placental explants (B) compared to wild type parasites. C) The number of host genes besides Ccl22 that are GRA28-dependent in placental explants is relatively small, suggesting that GRA28 is a highly specific inducer of Ccl22 in mouse placental tissue. D,E) Mouse serum (D) and peritoneal lavage (E) levels of Ccl22 48 h-post-infection are dependent on GRA28.

### GRA28-deficient parasites have distinct inflammatory and dissemination phenotypes in the acute and chronic phases of infection, respectively

To determine the impact of *T. gondii* GRA28 *in vivo* we indexed differences in mouse behavior relevant to inflammatory responses and quantified differences in infection-induced weight loss and total morbidity after infection of BALB/cJ mice with either RH:WT or RH*ΔGRA28 T. gondii*. We observed no significant differences in morbidity or weight loss (**Figure 7a,b**). However, when we scored (**Figure 7 Supplement 1**) mice over the course of infection as to the extent of inflammation-induced behavioral changes, we observed significantly heightened fur ruffling in the *ΔGRA28*-infected mice on days 6 and 7 post-infection (**Figure 7c**), despite the fact that mortality was unchanged.

**Figure 7.**
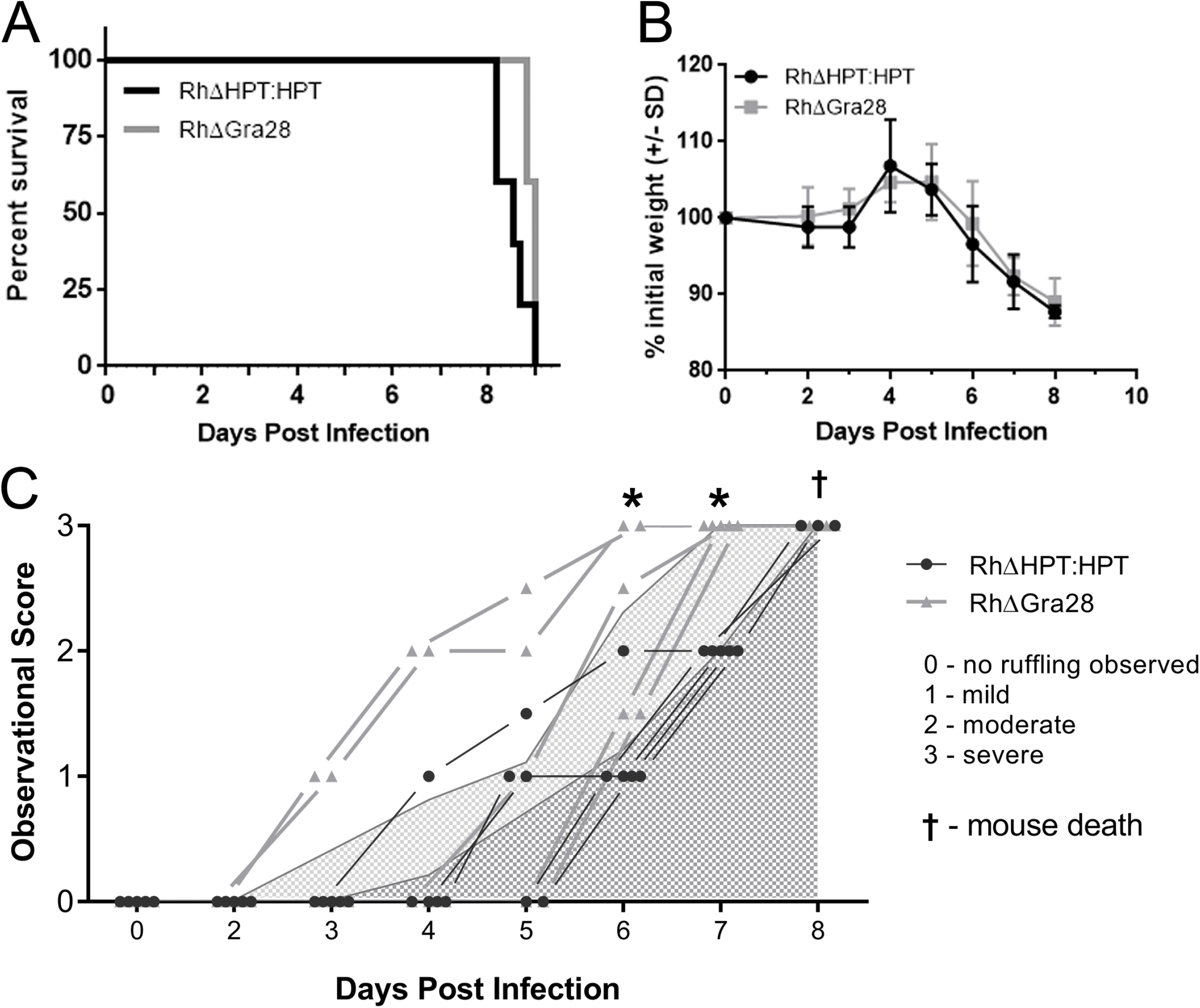
A) Mortality and B) weight loss does not significantly differ in mice infected with WT and ΔGRA28 type 1 strain *T. gondii* parasites. C) Behavioral changes and fur ruffling phenotypes associated with infection are exacerbated during the acute phase of infection for mice infected with ΔGRA28 parasites compared to WT based on a phenotype scoring system. Specifically, mice infected with ΔGRA28 parasites (light grey lines and triangles) exhibited infection-related symptoms at earlier time points compared to those infected with WT parasites (black lines and circles). Curves beneath lines are the average across all mice at that time point (with lighter grey representing Δ*GRA28* knockout parasites and darker grey representing wild type). *: P<0.05 after Two Way ANOVA and followed by multiple comparisons at each time point.

We also generated *ΔGRA28* parasites in a Type 2 *T. gondii* background that had been previously engineered to express luciferase and GFP (specifically ME49Δ*HPT:LUC*; (27, 28)) to permit non-invasive quantification of parasite burden and dissemination over the course of infection. For the ME49 strain infections we observed only minor and non-significant differences in mouse morbidity and weight loss (**Figure 8a,b**). However, during the acute phase of infection we observed slight differences in parasite burden between ME49Δ*HPT:LUC* (WT) and ME49*ΔGRA28*-infected mice, with burden being significantly *higher* in ME49*ΔGRA28* compared to WT on day 9 post-infection (**Figure 8c**). This difference was not due to experimental variation in parasite input between strains since parasite burden was indistinguishable during the first 6 days post-infection (**Figure 8c**). In contrast to these minor differences during the acute phase of infection, we observed more dramatic differences in parasite burden during the later stages of infection. Specifically, quantification of *in vivo* bioluminescence data taken dorsally on days 14 and 15 post-infection revealed that WT parasites were of much greater abundance in the brains compared to those infected with ME49*ΔGRA28* (**Figure 8d,e**).

**Figure 8:**
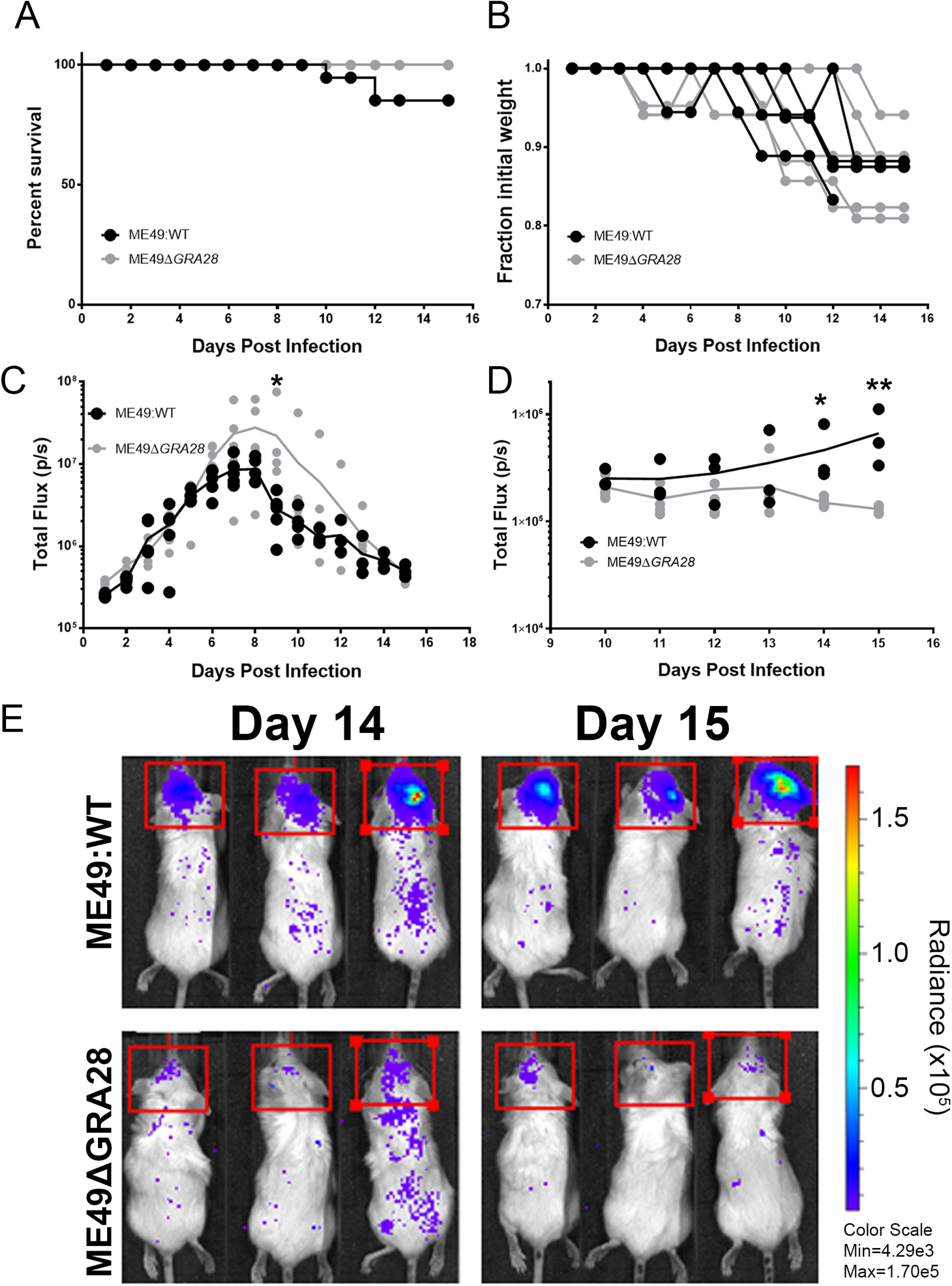
Impact of the *GRA28* gene on proliferation and dissemination of Type II parasites expressing luciferase. Neither A) mortality nor B) mouse weight loss were significantly different in mice infected with WT or *ΔGRA28 T. gondii* Type II (ME49) strain parasites. C) Throughout the acute phase of infection parasite-derived bioluminescence of WT and *ΔGRA28* parasites in the peritoneal cavity was measured by imaging the animals ventrally. Burden was similar (P>0.05) between parasite strains at all time points except for D9 post-infection, when signal was significantly higher in *ΔGRA28*-infected mice compared to WT mice (*P*=0.014). D,E) Starting on D10 post-infection we imaged mice both ventrally and dorsally to visualize dissemination to and proliferation in the mouse brain. In contrast to the acute phase we observed a consistently lower level of parasite-derived bioluminescence in the brains of mice infected with *ΔGRA28* parasites compared to WT parasites on days 14 and 15 post-infection (*P*=0.016 and 0.0002, respectively). All statistical tests were performed on log2-transformed bioluminescent data.

## DISCUSSION

*T. gondii*-infected host cells have dramatically altered transcriptomes compared to uninfected cells, and effectors that are secreted from the parasite during invasion drive most, but not all, of these changes (29). To date, the vast majority of these parasite effectors are derived from the dense granule and rhoptry organelles. We previously identified that *T. gondii* induces the production of CCL22 in human placental trophoblasts, while human foreskin fibroblasts do not exhibit this chemokine induction during *T. gondii* infection (4). Additionally, our previous work has shown that this induction required parasite invasion and the effector chaperone-like *T. gondii* gene product MYR1 (4, 13). While *T. gondii* induces CCL22 during infection of a variety of cell types from both mice and humans (18, 30), including at the transcriptional level in mouse brain (31), placental cell CCL22 induction is driven by a highly specific parasite effector, GRA28. CCL22 production is considered to be an indication of M2 macrophage polarization, and macrophage polarization has been linked to strain-specific *T. gondii* effectors like ROP16 and GRA15 (32). The impact of GRA28 is distinct from these effectors because CCL22 induction occurs similarly in all three canonical strain types, and GRA28 does not alter the expression of other M2-associated genes (such as *IL4*, *IL10*, *IL13* or *ARG1*), in the cell types that we have assayed. The specificity of GRA28 for only a few target genes is novel compared to effectors like GRA15 and ROP16 (24, 33) that alter the abundance of hundreds of transcripts.

Transcriptional co-regulation has been used in other systems as a means to identify members of protein complexes (34), but to our knowledge this is the first time this approach has been successfully applied at this scale in *T. gondii*. We used 396 microarray datasets derived from multiple *T. gondii* life stages and experimental manipulations to provide enough variation to better distinguish subclusters within closely-related gene families. Genes encoding dense granule proteins are among the most highly expressed in the *T. gondii* genome, making them more difficult to separate from one another, but they still clustered into two distinct groups with functional themes. The MYR1/GRA28 cluster harbored a handful of known secreted dense granule effectors, while the other contained genes encoding dense granule structural proteins or those that are secreted into the vacuole but do not traffic to the host cell. We anticipate that the former cluster can be exploited further to identify additional MYR1-trafficked, and putatively host-modulating, effectors while the latter has highlighted new candidates important in dense granule structure or function within the parasite. The entire dataset is available for download as a text file at Figshare (doi: 10.6084/m9.figshare.16451832) so that these data can be mined to identify candidates for membership in other critical *T. gondii*-specific protein complexes.

*GRA28* was previously shown to encode a dense granule protein secreted from the parasite into the host cell where it trafficked to the host cell nucleus (25), but its impact on the host cell was unknown. Its natural presence in the host nucleus during infection has also been further confirmed using proteomics, where it was found to be one of the more abundant *T. gondii* proteins in the nucleus of the infected host cell (35). The fact that it affects the abundance of only a small number of chemokine-encoding genes at the transcriptional level suggests that it modulates transcriptional activity via direct interactions with transcription factors and/or upstream regulatory sequences. Other *T. gondii* effectors traffic to the host nucleus but this is not always critical for function. For example, ROP16 localization to the host cell nucleus is dispensable for its primary function of phosphorylating STAT6 which occurs in the cytoplasm of the host cell (36). Other *T. gondii* effectors like IST (15, 37) and GRA24 (16) function within the host cell nucleus, but many of these mediate changes in hundreds of transcripts via their cooperation with existing transcriptional suppressors (IST; (15, 37)) or activators (GRA24; (16)). It remains to be seen if the function of GRA28 can occur independent of nuclear trafficking or if this ultimate localization is required for chemokine induction, but its specificity for downstream genes raises the interesting hypothesis that it may function directly, possibly as a heterologous transcription factor.

The signaling pathway governing GRA28 function is unknown but some clues can be found in our pathway analyses which suggest a role for GRA28 in mediating changes in key immunity-related host cell signaling pathways. The transcription factor genes *JUN*, *FOS* and components of the NFκB complex were consistently linked to the GRA28-dependent host transcripts. LPS is a well-known activator of both NFκB and C-Jun activity in THP-1 cells (38, 39), and this can occur via Toll-like receptor activation (40). However *T. gondii* induction of CCL22 was not fully dependent upon host MYD88, since MYD88^(-/-)^ THP-1 cells still produced significant amounts of CCL22 in response to *T. gondii* infection (**Figure 1 Supplement 1c**). The difference in CCL22 production by the MYD88^(-/-)^ cells in comparison to the WT cells should also be considered in light of the fact that the cell lines have different origins (and therefore distinct passage histories which could have the more subtle effects shown on CCL22 production after infection). A distinct cluster of GRA28-dependent host genes was identified that encoded gene products involved in proteoglycan synthesis, including the rate-limiting enzyme XYLT1 . *T. gondii* attachment to host cells is mediated by interactions between parasite adhesins and host cell surface sulfated proteoglycans (PG) like heparan sulfate (41, 42), and *T. gondii* adheres poorly to cells with genetically or enzymatically depleted levels of surface sulfated proteoglycans (41, 42). Therefore direct and/or indirect modulation of XYLT1 transcript levels by GRA28 may serve to make infected cells susceptible to adhesion, and ultimately invasion, by *T. gondii* or any other pathogens that depend on surface proteoglycans.

GRA28 had no impact on transcript levels of the gene encoding CCL17 which is commonly co-regulated along with *CCL22*. Mouse macrophages infected with *T. gondii* produce Ccl17 and Ccl22 and this is due, at least in part, to another *T. gondii* effector GRA18 (18). Using the same *GRA18* knockout lines (kindly provided by the Bougdour lab) we found that GRA18 had no impact on CCL22 production at the transcriptional (not shown) or protein (**Figure 4**) level in human THP-1 cells, suggesting that GRA18 and GRA28 have distinct targets. This is also consistent with the observation that Ccl22 induction in RAW macrophages is only partially dependent on GRA18 and β-Catenin signaling, in contrast to Ccl17 and Ccl24. Finally, in our work we used lower MOIs (2-3 here compared to 5-6 in (18)). Regardless, GRA28 appears to be the more potent modulator of Ccl22 production compared to GRA18, while Ccl17 appears to be much more dependent on GRA18. It is exciting to speculate that *T. gondii GRA28* has evolved to uniquely target CCL22 as a means to gain access to the fetal compartment since this chemokine is potently induced in placental cells and this chemokine plays a role in immune tolerance during pregnancy (7). However, as shown clearly in this study, GRA28 also alters monocyte/macrophage CCL22 production, making it equally plausible that this intricate molecular relationship developed first as a more generalized immune evasion (via suppression) strategy.

The role of specific chemokines like CCL22 during *T. gondii* infection is poorly understood but the discovery of *GRA28* allows this to be addressed more directly using *T. gondii ΔGRA28* parasites from different genetic backgrounds. Hypervirulent *T. gondii* RH strain *ΔGRA28* parasites caused inflammation-related behavioral changes earlier during infection in mice, compared to mice infected with WT parasites, suggesting that GRA28 functions to suppress inflammatory responses (likely due to induction of CCL22 although we did not test this directly). This could arise via GRA28-mediated recruitment and/or activation of regulatory T cells to the site of infection. These behavioral changes occurred without an effect on the acute virulence phenotype as all mice succumbed to the infection with similar kinetics, which is consistent with an impact of GRA28 on suppressing inflammatory responses without altering the ability of the mouse to control parasite replication. However, after infections using the type 1 parasite genotype we observed a significant reduction in *ΔGRA28* parasite burden in the brain compared to wild type parasites. This effect was unexpected given the fact that parasite burden was statistically equivalent during the acute phase of infection, but points to a potential important role for GRA28 in altering the host innate immune response in a manner that increases host susceptibility to dissemination of *T. gondii* across critical barriers like that guarding the CNS. *T. gondii* can infect blood-brain barrier epithelial cells as a means to cross into the host CNS (43), so GRA28 may promote parasite survival at this critical interface by recruitment of regulatory T cells or other cell types that might downregulate inflammatory responses.

**Summary:** Taken together our data point to a specific role of *T. gondii* GRA28 in modulating chemokine production in the infected cell. Importantly, this effect occurs only in certain cell types, including cells from both human and mouse placenta. A relatively small number of host chemokines are affected by parasites expressing this gene, and it plays a role in both modulation of the inflammatory response (as evidenced by mouse behavior and appearance during infection) and ultimately parasite dissemination to “privileged” sites like the CNS.

## METHODS

### Cell Culture

All cell and tissue cultures were incubated at 37°C and 5% CO_2_. All media were supplemented with 10% fetal bovine serum (FBS; Atlas Biologicals), 2 mM L-glutamine, and 50 mg/mL penicillin-streptomycin. Human foreskin fibroblast (HFF) cells were grown in Dulbecco’s Modified Eagle Medium (DMEM; Gibco), Raw264.7 cells were grown in DMEM (Gibco) with 10 mM HEPES, and THP-1 cells were grown in RPMI 1640 medium (Corning). THP-1 cells were assayed for viability using Trypan Blue staining (0.4%) (Gibco), counted, spun at 120 x g for 10 minutes at 24°C and medium replaced with supplemented DMEM prior to infection. All THP-1 cell numbers listed are based on trypan blue-negative cells.

### Human Placental Explants

Human placental tissue from less than 24 weeks of gestation was obtained, cultured, and infected with *T. gondii* as described previously (44).

### Mouse Placental Explants

Mouse placental tissues were obtained by dissection of E12.5 or 18.5 Swiss Webster mice. Upon removing the fetuses from the mother, the placentas were dissected away from other tissues and placed into pre-warmed 37°C PBS. The placentas were washed 3x in fresh pre-warmed PBS. Each placenta was then cut in half with sterilized surgical scissors and each half was placed into a well on a plate with pre-warmed 37°C DMEM with 10 mM HEPES, 10% FBS, 2 mM L-glutamine, and 50 mg/mL penicillin-streptomycin. Each placenta had one half-piece of tissue represented in each treatment group. For *T. gondii* infections, isolated tissue was infected immediately with 5.0 x 10^5^ – 2.0 x 10^6^ parasites for ∼24 hours.

### Parasites

Type 1 (RH, GT1), Type 2 (Me49, Pru), Type 3 (Veg, CEP) *Toxoplasma gondii* tachyzoites and sporozoites, *Neospora caninum* (NC-1) tachyzoites, and *Hammondia hammondi* (HhCatAmer and HhCatEth1; (45, 46)) sporozoites were used in this study. Sporozoites were excysted from sporulated oocysts as described (47) and either used immediately or grown for 24 h in human foreskin fibroblasts prior to being used in controlled infections. Tachyzoites were maintained by continual passage in human foreskin fibroblast (HFF) cultures incubated at 37°C and 5% CO_2_ in DMEM supplemented with 10% fetal bovine serum (FBS) (Atlas Biologicals), 2 mM L-glutamine, and 50 mg/mL penicillin-streptomycin. The Rh*YFP* strain was a gift from David Roos (University of Pennsylvania), the Rh*ΔMYR1* and Rh*ΔMYR1:MYR1_c_* parasites (13) were a gift from John Boothroyd (Stanford University), the PRUΔGRA18 and complemented knockout parasites were shared by Alexandre Bougdour (18), and the PruΔ*Toxofilin* KO parasites (19) were a gift from Melissa Lodoen (UC Irvine). For infections, infected monolayers were washed with fresh cDMEM and then scraped and syringe lysed to release tachyzoites. These tachyzoites were then passed through a 5 µM syringe filter and counted. Parasites were then centrifuged at 800 x g for 10 minutes at 24°C and resuspended and diluted in cDMEM before being used in infections. Mock treatments were produced by filtering the same parasites through a 0.22 µM syringe filter and exposing host cells to the same volume of the filtrate as was used for parasite infections. Freeze treatments were produced by subjecting the parasites to -80°C for 15 minutes, fixation treatments by exposure to 4% paraformaldehyde for 10 minutes followed by washing in PBS, and sonication treatments by sonicating at 0°C using five 30-second bursts at 50 amps with 30-second cooling intervals in between bursts followed by microcentrifugation at 800 x g for 10 minutes to generate soluble (S) and pellet (P) fractions.

### Invasion inhibitor assays

For Cytochalasin-D (Cyt-D) treatment, parasites were pre-treated with 10 µg/mL of Cyt-D in cDMEM for 1 hour and then used to infect cells in the presence of 10 µg/mL of Cyt-D in cDMEM for the duration of infection. Vehicle of Cyt-D is DMSO (40 µL per mL of cDMEM). For 4- Bromophenacyl bromide (4-BPB) treatment, parasites were pre-treated with either 0.5 or 1 µM of 4-BPB for 15 minutes. 4-BPB was dissolved directly in cDMEM. Parasites were then washed twice with normal cDMEM with 10-minute 800 x g spin steps between each wash, and then used to infect cells in the presence of normal cDMEM.

### Plaque Assays

Parasites were serially diluted in media and used to infect monolayers so that each tissue culture flask of HFFs received 100 parasites. These flasks were then incubated at 37°C in 5% CO_2_ undisturbed for 5-7 days. At the end of the incubation period, each flask was counted for number of plaques present and parasite viability was calculated. Crystal violet staining was used to count plaques as follows: the monolayer was washed with PBS and fixed for 5 minutes with ethanol. Then crystal violet solution (12.5 g crystal violet in 125 mL ethanol mixed with 500 mL 1% ammonium oxalate in water) was introduced to the monolayer and allowed to stain for 5 minutes. The monolayer was then washed extensively with PBS and allowed to air dry prior to counting plaques.

### Candidate gene identification using transcript level correlation analysis

To identify candidate effectors for inducing CCL22 we exploited the fact that the CCL22 induction response in THP-1 and placental cells required the presence of the *T. gondii* effector translocation protein MYR1 (13, 14). We hypothesized that MYR1-dependent effectors would have similar transcript abundance profiles across diverse expression datasets. We downloaded 396 publicly available *T. gondii* microarray expression datasets from the Gene Expression Omnibus platform hosted by the NCBI (48). We loaded and processed each CEL file using the “affy” module implemented in R (49). Data were processed and normalized using the following commands: bgcorrect.method = “rma”, normalize.method = “quantiles”, pmcorrect.method = “pmonly”, summary.method = “medianpolish”. RMA-normalized data were exported, re-imported into R and then transposed. An all-versus-all Pearson correlation matrix was generated using the “cor” function from the R:Stats base module. Probe names on the Affymetrix array (in the format of XX.mXXXXXX) were converted to current TGME49 gene models using data downloaded from ToxoDB and the Vlookup function in Microsoft Excel. In some cases the microarray annotations could not be matched to current TGME49 gene model names and are shown as blanks in plots. This correlation matrix, the normalized array data used to generate it, and a key to convert Affymetrix probe names current gene model names are all available on Figshare (10.6084/m9.figshare.16451832). To analyze this correlation matrix we used hierarchical clustering tools implemented in R including heatmap.2 (from the gplots package) and the dendextend package. To identify candidate genes using this matrix we calculated the mean correlations between 5 bait genes and all other queried genes from the microarray. The bait genes were known to encode either components of the MYR complex themselves or known TgMYR substrates (TgMYR1; (13), TgMYR2, TgMYR3 (14), TgGRA24 (16) and TgIST (15)). Most of the candidate CCL22-inducing genes were identified based on having a) an average correlation with the 5 “bait” genes listed above ≥0.7, b) a dN/dS ratio ≥2, c) and the presence of a predicted signal peptide OR at least one transmembrane domain (which we reasoned could be a cryptic signal peptide if the wrong start codon was chosen for the current gene annotation).

### De novo transcript assembly

To identify and assemble transcripts coding for GRA28 we used *de novo* transcript assembler Trinity ((50); version 2.6.6; default settings) using triplicate RNAseq datasets from WT *T. gondii* RH parasites infecting THP-1 cells (see below). Assembled transcripts with similarity to the predicted *GRA28* gene (TGME49_231960) were identified using BLASTN and BLASTX. Primary plots were generated using GenePalette software (51) and then modified.

### CRISPR-mediated gene disruption and validation of knockouts

The pSAG1::CAS9-U6::sgUPRT plasmid provided David Sibley (Addgene plasmid #54467;(52)) was modified using the Q5 Site-Directed Mutagenesis Kit (NEB) so that the gRNA sequence was replaced with two restriction enzyme sites (sequence: GTTTAAACGGCCGGCC) for PseI (NEB R0560S) and FseI (NEB R0588S). This modified plasmid was then used as the template for all future Q5 reactions. Two unique gRNA sequences were created for each candidate gene by utilizing the genomic sequences for *T. gondii* GT1 (toxodb.org) and E-CRISP (e-crisp.org) using the ToxoDB-7.1.31 reference genome. A forward primer for each gRNA was created for use with the modified pSAG plasmid, with the unique gRNA sequence followed by a section of the plasmid scaffolding (GTTTTAGAGCTAGAAATAGCAAG). The reverse primer used with this plasmid is AACTTGACATCCCCATTTAC. The gRNA sequences for the genes mentioned in this study and the primers used to validate the knockouts are listed in **Supplementary Table 1**.

A plasmid was created for each gene of interest (GOI) using the modified pSAG plasmid template and the Q5 Site-Directed Mutagenesis Kit (NEB) by following the manufacturer’s protocol with a few adaptations. The KLD enzyme step was extended to 60 minutes incubation at room temperature, and following the KLD enzyme step the product was heated to 65°C for 20 minutes and then double digested with PseI and FseI in CutSmart buffer (NEB) for 60 minutes at 37°C to remove any remaining plasmid that was not eliminated by the DpnI in the KLD step. This digested product was then heated to 65°C for 20 minutes to deactivate the enzymes prior to transformation, plasmid isolation and sequencing to validate insertion of the correct gRNA sequence (pSAG:GOI:gRNA). Parasites were transfected with either a single gRNA plasmid or equal amounts of plasmids encoding two gRNAs targeting the same gene. For validation of knockouts after cloning, a clone was considered a knockout if PCR across a targeted cut site failed or if a PCR reaction across the entire gene (just upstream and downstream of the start and stop codons, respectively) failed. In some cases where all PCR reactions worked (indicating that the plasmid failed to insert at the gRNA target site), the amplified band was sequenced and a clone was considered a knockout if insertions/deletions were identified near the gRNA binding that resulted in frame shifts and premature stop codons.

### Parasite transfections

In general, transfections were performed using standard approaches. Briefly, parasite suspensions were obtained by needle passage (25 and 27 gauge needles) and then pelleted for 10 minutes at 800 x g. Parasites (∼2 x 10^7^ per transfection) were re-suspended in Cytomix (120 mM KCl; 0.15 mM CaCl_2_; 10 mM KPO_4_; 25 mM Hepes, 2mM EDTA, 5mM MgCl_2_; pH to 7.6) containing GSH and ATP and electroporated at 1.6 Kv with a capacitance setting of 25 µF using a BTX ECM600 Electroporator. Transfected parasites were then used to infect coverslips and/or flasks of confluent HFFs and placed under appropriate selection. For candidate gene knockouts, ∼2×10^7^ *T. gondii* RHΔHPT parasites were transfected with ∼30-50 µg of the relevant pSAG:GOI:gRNA plasmid(s) (described above) along with 2-5 µg of an empty pGRA-HA-HPT (53) plasmid. Parasites were placed under selection the next day and cloned by limiting dilution after 2-3 passages. Individual clones were screened for gene deletion by PCR and sequencing to permit identification of both target gene disruptions (via insertion of the pGRA-HA-HPT plasmid at the CAS9 cut site) or mutation via DNA repair events at the CAS9 cut site. For HA-tagging experiments, Type 1 (RH) *GRA28* exon 1 (residues 1-498) was C-terminally HA tagged by cloning into the *T. gondii* expression plasmid pGRA-HA-HPT (53). This plasmid drives protein expression using the highly active *GRA1* promoter. TgRHΔ*HPT* or *N. caninum* Liverpool (NcLIVΔ*HPT*; (54)) parasites were transfected with ∼40-60 µg of *GRA28* exon 1 plasmid and cells were grown overnight in normal media. For analysis of transiently transfected parasites, cells were only grown for 18 h post-transfection while for stable transfection parasites were grown for 2-3 passages in media containing 50 µg/mL of mycophenolic acid and xanthine. Cells were fixed with 4% PFA and permeabilized in 0.1%Triton/PBS. Samples were probed with anti-HA rat monoclonal antibody (3F10 clone, Roche) diluted to 0.1 mg/mL in 0.1%Triton-PBS buffer and washed four times in PBS. Samples were then incubated in 488 goat anti-rat (Life Technologies Alexa Fluor H+L) followed by PBS washes. All samples were mounted in Vectashield with DAPI (Vector laboratories). For genetic complementation, TgRH*ΔGRA28* parasites were transfected the exon 1 construct used above or a construct amplified from genomic DNA encompassing the start and stop codons of the GT1 version of GRA28 (TGGT1_231960). Expression plasmids (∼30 µg) were co-transfected along ∼5 µg of pLIC_3xHA_DHFR* plasmid (kindly provided by Vern Carruthers; (55)), and populations were placed under 1 µM pyrimethamine selection for 2-3 weeks, and then used to infect THP-1 cells for 24 h as above followed by assays to quantify CCL22 in culture supernatants by ELISA.

### Fluorescence image analysis

To compare signal intensity in the nucleus of host cells infected with either *T. gondii* or *N. caninum* parasites transiently transfected with the HA-tagged GRA28 exon 1 construct, we scanned stained coverslips for GRA28-HA-positive vacuoles, and then used Fiji (an implementation of Imagej;) to calculate 1) the average HA signal intensity in the nucleus of the infected host cell (AvgIntInf), 2) the average HA signal intensity in the nucleus of a neighboring, uninfected host cell (AvgIntUninf) and 3) the average signal intensity of the parasite-containing vacuole (AvgIntVacuole). We then used the following calculation to determine the normalized, background-subtracted nuclear signaling intensity:

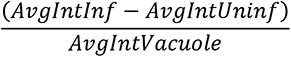

Example images of this process are shown in **Figure 5** and **Figure 5 Supplement 1**. Data were log10 transformed prior to performing a Student’s T-test.

### RNA-seq

RNA was isolated from cultures using the RNeasy Mini Kit (QIAGEN) and its associated RNase-Free DNase digestion set (QIAGEN), following the manufacturer’s protocol for mammalian cells. An Agilent Bioanalyzer was used to check the quality of the RNA samples. Tru-Seq stranded mRNA libraries were generated from 5-17 ng/µl of mRNA for THP-1 cells, and 50-120 ng/µL of mRNA for murine PECs and placental explants, and sequenced with an Illumina NextSeq 500 sequencer. mRNA-Seq FASTQ reads were mapped to the human reference genome (Homo sapiens v81; hg38) using default options on CLC Genomics Workbench 11 (Qiagen). Total gene reads (with at least 1 read count) were exported from CLC Genomics Workbench and used for DESeq2 (56) to perform differential expression analysis using methods outlined previously (e.g., (4)). Data were evaluated using principal component analysis (embedded in the DESeq2 package) and genes were deemed to be significantly expressed if the log_2_ fold-change was ≥ 1 or ≤ -1 and with a P_adj_ value <0.01. Gene set enrichment analysis (GSEA;(57)) and Ingenuity Pathway Analysis (QIAGEN; (58)) software were used to compare gene sets that were differentially regulated after infection with WT and Δ*GRA28* parasites.

### CCL22 and Ccl22 ELISA

CCL22/Ccl22 ELISAs were performed on culture supernatants (undiluted or diluted when necessary) using Immulon 4HBX flat bottom microtiter plates with the human CCL22/MDC DuoSet ELISA (R&D Systems DY336) or the mouse Ccl22/Mdc DuoSet ELISA (R&D Systems DY439) per the manufacturer’s instructions.

### Mouse Experiments with WT and *ΔGRA28* parasites

To determine the impact of *T. gondii* candidate effectors on mouse morbidity and cytokine production, BALB/cJ mice from Jackson laboratories (4-6 week old, female) were injected intraperitoneally with 200 µL of PBS containing 1.0 x 10^6^ *T. gondii* tachyzoites, or 200 µL of 0.22 µM filtered parasite solution as a mock treatment. Mice were sacrificed at 48 hours post infection (HPI), and peritoneal exudate cells (PECs) were collected by injecting 3 mL sterile PBS into the abdominal cavity, rocking the mouse to mix the PBS, and siphoning the PBS solution into a sterile conical tube. The solution was then centrifuged at 1,000 x g for 10 minutes at 24°C, the supernatant was collected, and RNA was extracted from the pellet for RT-qPCR and RNA-seq. Blood was collected in Sarstedt Microvette CB 300 Z tubes by cardiac puncture and centrifuged at 10,000 x g for 5 minutes to separate the serum. Mice were infected with RHΔ*HPT* (wild type), RH*ΔMYR1* or RH*ΔGRA28* parasites depending on the experiment.

To determine the impact of *GRA28* deletion on *T. gondii* proliferation and dissemination, female BALB/cJ mice (4 weeks old) from Jackson laboratories were injected intraperitoneally with 200 μL PBS containing 100 *T. gondii* tachyzoites. In one experiment five mice received an injection of RH*ΔGRA28* parasites, while the other five received an injection of the transfection control parasite that was transfected with empty vector (RhΔ*HPT:HPT*). For behavioral indices of inflammatory responses, photographs of the mice were taken dorsally and laterally every 4-6 hours for the entire duration of the infection. Mice were visually scored 0-3 based on the presence of fur ruffling, the location of ruffling, and the presence of skin redness/irritation (**Figure 7 Supplement 1**). **0)** No fur ruffling or red/irritated skin present. **1)** Mild ruffling present predominantly located on the head and back of the neck. No red/irritated skin visible. **2)** Moderate ruffling present - fur forms larger clumps and extends to the rest of the body. Skin may be visible through the clumps but is not red or irritated. **3)** Severe ruffling characterized by fur ruffling across the entire body with visibly red/irritated skin in between fur clumps.

In a second experiment mice were infected with 1000 tachyzoites of *T. gondii* strain ME49:*LUCΔGRA28* or a passage-matched wild type strain (27, 28). Mice were imaged daily after injection of D-Luciferin as described previously (28, 59) using an IVIS Lumina II *in vivo* bioluminescence imaging system (with ventral imaging occurring on all days post-infection and ventral and dorsal imaging occurring starting on day 10 post-infection). Animals were anaesthetized using 2% isoflurane during the 4-8 minute imaging period (ventrally and dorsally where applicable). When necessary, blood was collected via submandibular lancet puncture, collected into Sarstedt Microvette CB 300 Z tubes and spun at 10,000 x g for 5 minutes to separate the serum. Mice were monitored extensively over the course of infection for symptoms of morbidity and humanely euthanized. All animal procedures were approved by the Division of Laboratory Animal Resources and IACUC and our animal facilities are routinely inspected by the USDA and local IACUC committee.

### Statistics

All statistics were performed in Graphpad Prism for Windows (versions 7 or 9; GraphPad Software, La Jolla, CA). For most two treatment assays, we used unpaired, two-tailed student’s T test, and for multi-treatment/condition experiments we used one- or two-way ANOVA followed by multiple comparisons post-hoc tests. Individual comparisons are listed for each assay in the text and figure legend, and only pre-planned comparisons were performed to minimize Type 1 error. *In vivo* bioluminescence data (total flux; photons/s) and nuclear staining intensity data (comparing nuclear trafficking of *T. gondii* GRA28 when expressed in *T. gondii* or *N. caninum*) were log_10_-transformed prior to statistical analysis.

## Acknowledgements

The authors would like to thank Dr. Alexandre Bougdour (Inserm; Grenoble, France) for providing GRA18 knockout and complemented strains, Drs. John Boothroyd and Michael Panas (Stanford University) for helpful discussions and providing MYR1 knockout and complemented strains, and Dr. Peter Bradley (University of California-Los Angeles) for helpful discussions and sharing reagents related to GRA28 that facilitated this work.

## SUPPLEMENTAL FIGURE LEGENDS

**Figure 1 Supplement 1:** A) Type 1 (Rh88), Type 2 (Pru) and Type 3 (CTG) *T. gondii* strains all induce CCL22 in THP-1 cells. For mock cells were treated with 0.2 micron-filtered parasite suspensions. B) *H. hammondi* induces CCL22 production in THP-1 cells while *N. caninum* does not when compared to mock-treated cells. C) The *T. gondii* gene MYR1 is required for CCL22 production by THP-1 cells after infection. *T. gondii* ΔMYR1 parasites were compared to ΔMYR1:MYR1-complemented parasites and only the MYR1-complemented parasites induced secretion of CCL22 from wild type (black data points; left) or MYD88-knockout (blue data points; right) THP-1 cells. While MYD88 knockout cells produced significantly less CCL22 in response to MYR1-complemented parasites, they still produced CCL22 at levels much greater than ΔMYR1-infected cells or mock-treated cells. MYR1-knockout and complemented parasites were kindly provided by John Boothroyd and Michael Panas, Stanford University.

**Figure 2 Supplement 1:** Clusters of co-regulated genes share developmental expression profiles and functional activities. A) Transcript abundance correlation analysis (left) and clustered transcript abundance analysis (in RMA log2-normalized units) for 21 genes with transcripts known to increase in abundance during the tachyzoite to bradyzoite transition, including BAG1, LDH2 and enolase. Bar across the top of the expression heatmap indicates the life cycle stage source for each of the samples. Pie chart indicates that 19 of the 21 transcripts increase in abundance during pH-induced bradyzoite development according to the Bradyzoite Differentiation (3-day Time Series) dataset on ToxoDB.org. B) Cluster containing multiple ribosomal protein coding genes (for both the large and small subunits) showing high transcript abundance in tachyzoites and bradyzoites but comparatively low transcript abundance in samples taken from sporozoites and merozoites.

**Figure 3 Supplementary Figure 1:** Transcript level correlation analysis with the 6 bait genes used in this study (A) and clustered, normalized gene expression data (B) for all genes in the *T. gondii* genome annotated with the term “dense granule” in the product name or user comments. Dense granule protein coding genes fall into two major clusters in the correlation analysis, with the top cluster containing the known secreted effectors GRA15, GRA24, TgIST, GRA7 and GRA28.

**Figure 3 Supplementary Figure 2:** Validation of knockouts generated for the present study (A-D) of for GRA18 which was generated in another study (E; He *et al.*, eLife 2018;7:e39887). Clones that were validated as knockouts and used in the CCL22-induction assays shown in Figure 3b are labled with ***bold-italic font***. We validated using two paralell approaches for most of the knockouts: (1) amplifying across the site targeted by the protospacer(s) encoded by the transfected gRNA-expressing plasmid(s) where no amplification indicated a potential insertion of plasmid sequence into that location and (2) amplifying across the entire coding sequence of the gene, where no amplification also suggests insertion of the selectable marker and other plasmid sequence at at least one of the protospacer sites. In some cases multiple protospacer encoding plasmids were transfected into the same parasite population (and protospacer numbers are our own internal nomenclature), and in these cases it was possible to have insertion/disruption at both protospacer sites (and this ocurred in some instances). Primer sequences and gRNA target sites are shown in **Table S1**. A) Validation of four TGGT1_201390 knockout clones generated by batch transfection with gRNAs targeting two distinct sites in the TGGT1_201390 coding sequence (2 and 12). Left gel: Two primer sets (A and B) were used to amplify across the site targeted by the protospacer, and MAF1b primers were used as a positive control. Clones D11 and G11 likely had insertions at protospacer site 2 (as evidenced by the lack of amplification with primer set A), while clone D2 likely had an insertion in protospacer site 12 as evidenced by a lack of amplification with primer set B. Clone F2 had amplification of the correct size at both protospacer sites (2 and 12), but we could not amplify the coding sequence from the F2 clone (right gel, lane labeled F2), suggesting that the locus was disrupted in this strain as well. All four of these knockout clones were assayed in biological triplicate for CCL22 induction in THP-1 cells (Figure 3b). B) Validation of 3 GRA4 knockout clones out of 5 tested. Clones A1:D6 and B2:C1 (where A1 and B1 indicate the parasites were from distinct transfections) had a likely insertion in the GRA4 gene at protospacer site 0, while clone B2:B11 had a likely insertion at both sites (0 and 26). C) Validation of 4 GRA8 knockout clones. Parasites were transfected with a single gRNA expressing plasmid (targeting gRNA sequence 4) and PCR on all 4 clones failed to amplify across the gRNA 4 target site. All amplifications across the gRNA target site 27 worked as did the positive control amplification of MAF1b. D) Validation of 4 GRA28 knockout clones. Parasites were co-transfected with plasmids encoding gRNAs targeting sites 5 and 26 and queried using primers targeting the entire locus (sets A and B in this case), or flanking gRNA target sites 5 and 26 (sets C and D). PCR across the entire GRA28 locus for clones 1A4, 1D4 and 1D3 all failed to generate PCR products, suggesting that these 3 clones had disruptions in the *GRA28* gene. PCR on clone F3 with primer sets A and B gave a product of the expected size. PCR across the gRNA target site 26 failed for clone 1A4, while PCR across gRNA target site 5 failed for clones 1D4 and 1D3. Taken together these data indicate that clones 1A4, 1D4 and 1D3 were all GRA28 knockouts via insertion of selectable marker and accompanying plasmid sequences at the targeted gRNA sites. For clone 1F3 the locus seemed to be intact, but when we sequenced PCR products similar to those amplified by primer sets C and D (at gRNA target sits 5 and 26, respectively) we determined that clone 1F3 had a single base pair deletion at base 250 relative to the start codon of GRA28 which introduced a stop codon 100 bp downstream of the indel (as well as multiple stop codons in frame further downstream). Sequences across the gRNA 26 target site were identical to wild type. The deletion near gRNA target site 5 was within the gRNA protospacer sequence itself, just proximal to the PAM site (GTTCCGCTGGTGCCTT**C**ACC [TGG] was mutated to GTTCCGCTGGTGCCTT**_**ACC [TGG]). Therefore we treated 1F3 as a GRA28-null parasite strain. E) Validation of GRA18 knockouts received from the Bougdour lab: We received GRA18 knockout and wild type clones from the laboratory of Alexandre Bougdour and validated them using PCR. In this case we generated primers to amplify across the entire GRA18 locus, which was completely deleted using double homologous recombination with large sequences flanking the entire coding region. As expected we could amplify across the entire GRA18 locus in PRU:WT and GRA18-complemented strains, but failed to do so in the GRA18 knockout.

**Figure 4 Supplementary Figure 1:** Ingenuity pathway analysis of THP-1 cells infected with RH:WT or RH*ΔGRA28 T. gondii* identifies candidate upstream regulatory gene products that may be driving GRA28-dependent differences in transcript abundance. A) Genes of higher abundance in RH:WT-infected THP-1 cells were found to be significantly associated with multiple immunity-related regulatory genes, including those related to the NFκB pathway (e.g., NFκBIA, NFκB1 and REL). B) Hierarchical cluster of correlations in the amount of target gene overlap for each of the upstream regulators shown in A. A small cluster of the most highly correlated genes contained multiple genes relevant to NFkB activation (outlined in dotted green). C) As in B, most of the upstream regulators had the same downstream targets, and this was most evident for the AP-1 transcription factor complex (encoded by *FOS* and *JUN* genes) as well as *IL1B* and *ICAM1*. A heat map indicating fold-differences between RH:WT and RH*ΔGRA28*-infected THP-1 cells for these downstream targets is also shown. D) Heatmap showing transcript abundance for all transcriptional regulators that were found to be significantly altered in infected THP-1 cells in a GRA28-dependent manner (P<0.001; log_2_(fold-difference)≥1).

**Figure 4 Supplementary Figure 2:** C-JUN protein levels are induced by *T. gondii* infection in THP-1 cells but do not depend on the presence or GRA28 in the infecting strain. THP-1 cells were infected with the indicated strains (or mock-treated by exposing them to the same WT parasite suspensions as for infection but after sterile-filtering with a 0.2 µm filter) for 24 h and then C-JUN protein level was quantiifed using western blotting. Histone H3 levels served as a control and densitometry was used to calculate the C-JUN/Histone H3 ratio as a proxy for normalized C-JUN abundance. Two replicates, with an N=3 wells of cells for infections and N=2 wells for mock, are shown, each having similar results.

**Figure 5 Supplementary Figure 1:** A) Quantification of nuclear localization in *T. gondii* (N=5) and *N. caninum* (N=3) vacuoles expressing Exon 1 of *T. gondii* HA-tagged GRA28. Data were normalized for each image and HA-positive vacuole by subtracting mean HA-intensity in a nucleus neighboring the infected cell from the mean GFP intensity in the nucleus of the infected cell, and then dividing that by the mean HA-intensity of the parasite-containing vacuole. Expression in *T. gondii* led to significantly (P<0.01; T-test on the log10-normalized data) higher normalized intensity in the nucleus compared to when GRA28 was expressed in *N. caninum*. B,C) Schematic illustrating how data in A were collected and calculated for *T. gondii* (B) and *N. caninum* (C).

**Figure 7 Supplementary Figure 1**: Visual representation of pathology index scores. All images are the same individual from two viewpoints (lateral and dorsal) at four different timepoints of infection. **0)** No fur ruffling or red/irritated skin present. **1)** Mild ruffling present predominantly located on the head and back of the neck. No red/irritated skin visible. **2)** Moderate ruffling present - fur forms larger clumps and extends to the rest of the body. Skin may be visible through the clumps but is not red or irritated. **3)** Severe ruffling is characterized by ruffling across the entire body with visibly red/irritated skin.

